# Neuromodulation via muscarinic acetylcholine pathway can facilitate distinct, complementary, and sequential roles for NREM and REM states during sleep-dependent memory consolidation

**DOI:** 10.1101/2023.05.19.541465

**Authors:** Michael Satchell, Blaine Fry, Zahraa Noureddine, Alexis Simmons, Nicolette N. Ognjanovski, Sara J. Aton, Michal R. Zochowski

## Abstract

Across vertebrate species, sleep consists of repeating cycles of NREM followed by REM. However, the respective functions of NREM, REM, and their stereotypic cycling pattern are not well understood. Using a simplified biophysical network model, we show that NREM and REM sleep can play differential and critical roles in memory consolidation primarily regulated, based on state-specific changes in cholinergic signaling. Within this network, decreasing and increasing muscarinic acetylcholine (ACh) signaling during bouts of NREM and REM, respectively, differentially alters neuronal excitability and excitatory/inhibitory balance. During NREM, deactivation of inhibitory neurons leads to network-wide disinhibition and bursts of synchronized activity led by firing in engram neurons. These features strengthen connections from the original engram neurons to less-active network neurons. In contrast, during REM, an increase in network inhibition suppresses firing in all but the most-active excitatory neurons, leading to competitive strengthening/pruning of the memory trace. We tested the predictions of the model against *in vivo* recordings from mouse hippocampus during active sleep-dependent memory storage. Consistent with modeling results, we find that functional connectivity between CA1 neurons changes differentially at transition from NREM to REM sleep during learning. Returning to the model, we find that an iterative sequence of state-specific activations during NREM/REM cycling is essential for memory storage in the network, serving a critical role during simultaneous consolidation of multiple memories. Together these results provide a testable mechanistic hypothesis for the respective roles of NREM and REM sleep, and their universal relative timing, in memory consolidation.

**Significance statement:** Using a simplified computational model and *in vivo* recordings from mouse hippocampus, we show that NREM and REM sleep can play differential roles in memory consolidation. The specific neurophysiological features of the two sleep states allow for expansion of memory traces (during NREM) and prevention of overlap between different memory traces (during REM). These features are likely essential in the context of storing more than one new memory simultaneously within a brain network.

## Introduction

In vertebrate species, sequential cycling from wake to non-rapid eye movement (NREM) sleep to REM sleep is a universal pattern (1). The evolutionary origins of the physiologically-distinct NREM and REM states are a matter of speculation (2). However, available data suggest that both are essential for brain functions, including long-term memory consolidation and learning-associated synaptic plasticity (3-6). The two states have dramatically different features – including characteristic neuronal firing patterns and rate changes, network oscillations, and neuromodulation (3-5, 7-12) – suggesting NREM and REM likely have distinct effects on brain circuits during memory storage. Recent studies have identified molecular (13, 14) and electrophysiological (15-25) changes in neural circuits during post-learning NREM and REM sleep. A physiological variable which diverges widely between wake, NREM sleep, and REM sleep is the level of acetylcholine (ACh) release in the forebrain (26). ACh regulates neural excitability, and is essential for brain processes ranging from sleep-wake regulation to sensory detection (27-30). Cholinergic projections are found throughout the forebrain, midbrain and brainstem (31).

In the hippocampus, cholinergic inputs originating in the medial septum (MS)(32, 33) provide direct input to both principal neurons and interneurons (34). The septo-hippocampal projection is a critical regulator of hippocampal function, regulating theta oscillations vital in memory acquisition and storage (35). ACh acts through nicotinic and muscarinic receptors (36, 37). In the hippocampus, the most abundant nicotinic receptor subtypes are α7 and α4β2 receptors, expressed by pyramidal neurons and GABAergic interneurons (37-40). At the same time, ACh acts on five subtypes of muscarinic receptors, M1 through M5 (41). M1 muscarinic receptors, expressed on neuronal dendrites or somas, are the most abundant subtype in the hippocampus (42, 43). These receptors regulate the excitability of hippocampal neurons (44, 45) via coupling to slow non-inactivating potassium channels. Regulation of these channels’ corresponding ionic current (m-current), blocked when ACh is high, leads to a switch between Type 2 excitability (when concentration ACh is low) and Type 1 excitability (when concentration Ach is high) (46-49). Shifts from Type 1 to Type 2 excitability cause dramatic changes in firing frequency responses to injected current (i.e., f-I curve or gain function) (49). The f-I curve of Type 1 neurons is continuous and has a relatively large firing frequency response to incremental changes in excitatory current (49). Type 2 neurons have a discontinuous frequency increase from quiescence and exhibit relatively small frequency responses to changes in excitatory current (49). A concurrent change at the switch between Type 1 and Type 2 dynamics is a differential response of spike timing (i.e., an advance or delay in spiking) in response to weak excitatory input – the so-called phase response curve (PRC) (46); A Type 1 PRC is uniformly positive, meaning that excitatory synaptic inputs always advance the timing of the next spike. Type 2 dynamics are characterized by a biphasic PRC, with advances or delays possible (46, 49). This biphasic nature of the Type 2 PRCs mediates spike synchrony among neurons in a network (49, 50). Thus, Ach modulation at M1 receptors can dramatically change neurons’ frequency responses to excitatory input, as well as their capacity to synchronize their firing to one another.

At the network level, ACh may also modulate the relative activity levels of interneurons vs. principal neurons, to alter excitatory-inhibitory balance in brain networks (51, 52). In the hippocampus, ACh has the ability to selectively activate somatostatin-expressing (SST+) interneurons (53-55), which can affect the encoding, storage, and recall of memories. Inhibitory gating of activity in the DG network by SST+ interneurons in the hours following learning constrains hippocampal memory consolidation (56). Chemogenetic suppression of medial septal ACh projections to hippocampus after single-trial fear conditioning improves sleep-dependent fear memory consolidation and increases DG granule cell activity. Conversely, chemogenetic activation of ACh inputs suppresses DG granule cell activity and impairs memory consolidation (56). These results are consistent with prior research indicating that suppression of cholinergic signaling after learning benefits sleep-dependent, hippocampally-mediated memory consolidation in human subjects (57, 58).

Together, these findings suggest that NREM-associated suppression of ACh input to hippocampus is an essential feature for sleep-dependent memory consolidation. However, there are accumulating data suggesting that REM – a high-ACh brain state - also plays an important role in consolidation of hippocampus-dependent memories (59-62). Yet it remains unclear whether NREM and REM play overlapping, distinct, or complementary roles in the storage of information in neural networks. How the bidirectional changes in cholinergic signaling during these two states affect the dynamics and function of hippocampus is largely unknown.

Here, using *in silico* models, we investigate ACh-regulated mechanisms underlying differential NREM- and REM-specific effects on memory storage in neural networks. We show that disinhibition and excitability changes in principal cells, mediated by activation of m-current in response to reduced ACh signaling during NREM, increases recruitment of new neurons into newly-encoded memories’ engrams. In contrast, increased activation of inhibitory interneurons when m-current is blocked by ACh signaling during REM leads to competitive and selective pruning of neural representations associated with given memories. This REM-dependent mechanism is specifically important when multiple memories are to be consolidated simultaneously, as it naturally reduces overlap between different memory representations following a period of NREM. Repeated iterations of the NREM→REM cycle (such as those occurring across a night of sleep) lead to progressive expansion and segregation of memory traces in the network. Critically, reversal of the NREM→REM sequence (i.e., REM→NREM) led to failure of memory trace expansion. Together these data suggest that NREM and REM could play distinct and complementary roles in memory storage in the hippocampal network, and that the NREM→REM state sequence, which is so universal across various animal species, is an essential feature for memory consolidation.

## Results

### ACh-regulated neural network states differentially recruit surrounding neuron populations into a newly-formed engram

We used a reduced network to model ACh-dependent effects of NREM and REM sleep on previously-encoded memories. Based on experimental data (63), we constructed a reduced network composed of three cell populations. Within the model, initial memory traces (or”engram backbones”) are formed with a limited number of engram cells (i.e., excitatory neurons that are highly active, as is true after learning)(63). This highly-active backbone population comprises the first of the three cell populations. The second is composed of non-backbone excitatory neurons that are less excitable (LE) due to weaker excitatory input and greater input from inhibitory interneurons (64-68). The excitatory synapses impinging on these neurons (both connections from backbone neurons and reciprocal connections within the LE population) undergo spike timing dependent synaptic plasticity (STDP). The two excitatory networks have sparse (∼10%), largely random connectivity, with fast AMPA receptor kinetics (see **Methods**). Finally, the third cell population within the network is an interneuron population, which inhibits both backbone neurons and the LE population. This interneuron population is numerically smaller, but with a high connectivity density (50%). Consistent with available data (64-68), these interneurons differentially target the backbone and LE populations, providing a higher level of inhibition to LE neurons via both fast (GABA_A_-mediated) and slow (GABA_B_-mediated) receptor kinetics (see **Methods**). All three populations exhibit simultaneous, state-dependent changes in ACh signaling, modeled as a change in m-current conductance. Low concentrations of Ach present during NREM sleep are modeled as high m-current conductance 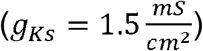, while high ACh concentrations present during REM sleep are modeled as low m-current conductance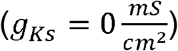.

We first investigated population activation patterns as a function of m-current conductance (**Fig. 1**) for a network with a single encoded memory (i.e., engram backbone [EB] population; **Fig. 1A**). Mean firing frequency changes for neurons within each population (i.e., backbone neurons, LE neurons and inhibitory interneurons) was measured as *g*_*Ks*_ was varied from 0 to 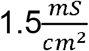. When ACh signaling was high (*g*_*ks*_= 0; e.g., during REM) both backbone neurons and inhibitory interneurons exhibited highly synchronous periodic activation at theta frequency (**Fig. 1B (bottom), C (right panel)**), driven by slow interneuron-driven inhibitory current targeting the backbone neurons. This mechanism is like that underlying pyramidal interneuron gamma (PING)(69, 70), but causes oscillatory modulation at a lower frequency determined by slow inhibitory receptor kinetics. The activation of excitatory backbone neurons generated a burst of firing among the interneurons, which in turn synchronously inhibited excitatory population until the slow inhibitory postsynaptic current (IPSC) decays, allowing the backbone neurons to subsequently fire again. Since the LE neurons were more strongly inhibited by the interneurons, they did not activate during that process (**Fig. 1D**), and their firing pattern is driven by random noise in the network (see **Methods**). Hence, the population dynamics resulted in strong spectral power in the theta band for the whole network, driven primarily by the activation of backbone cells and inhibitory interneurons (**Fig. 1C, right**).

**Figure 1.**
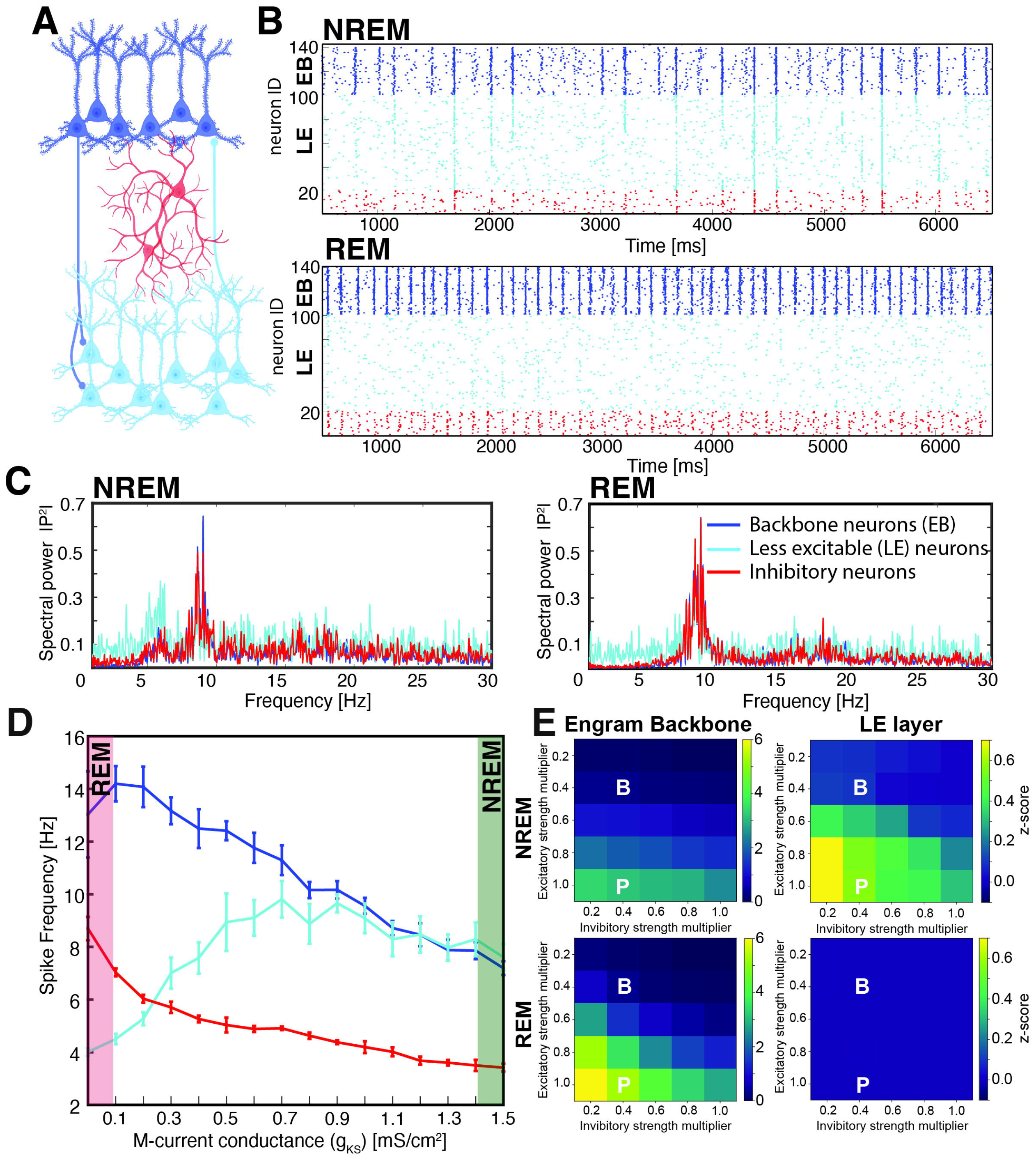
ACh-regulated current modulates activation of neuron populations during NREM-like and REM-like network states. ***A)*** Schematic of the model network, which consists of an excitatory, memory-encoding engram”backbone” (EB) neuron population (***dark blue***), inhibitory interneurons (***red***), and an excitatory population that is less excitable (LE) and non-memory-encoding (***teal***). ***B)*** Representative raster plot of firing patterns for the three populations during NREM (low ACh, high g_Ks_; **top**) and REM (high ACh, low g_Ks_; **bottom**). ***C)*** Spectral analysis of firing patterns during NREM (**top**) and REM (**bottom**) states. Data are shown for firing of the entire network (**left, *black line***) and separately for each neuron population (***colors denoted as above***). ***D)*** Mean firing frequency for the three network populations as a function of maximal g_Ks_ conductance. High g_Ks_ conductance during NREM (highlighted in ***green***) causes disinhibition of the LE population, and permits their participation in bursts driven by backbone neurons. Low g_Ks_ conductance during REM (highlighted in ***red***) increases activity in backbone and inhibitory neurons, while LE neuron activity is suppressed. Values indicate mean ± SEM, for 4 simulation runs. **E)** Mean functional connectivity between pairs of engram backbone (EB) neurons (**left**) and between pairs of LE neurons (**right**), as a function of strength of backbone neurons’ connections to both other excitatory neurons and inhibitory neurons. The default connection strength (see **Methods**) was multiplied by a factor as indicated on Y and X axis, respectively. Mean functional connectivity varies more with variation in synaptic strength in REM vs. NREM.

An increase in m-current conductance (due to decreased ACh) flattens the i-f curve of individual neurons, lowering their firing frequency response for a given input current (49, 71). When increasing *g*_*Ks*_, we observed heterogeneous responses among cell populations in the network, with the firing frequency of both inhibitory interneurons and backbone neurons (**Fig. 1D, dark blue and red lines**) decreasing. Lowering firing rates among inhibitory interneurons resulted in disinhibition of network LE neurons and an increase in their firing rate (**Fig. 1D, teal line**). Thus, during NREM, when *g*_*Ks*_ is high, LE neurons were more easily recruited into synchronous bursts, which emerged from type 2 neural excitability and biphasic PRCs (associated with high *g*_*Ks*_) (69, 70) with a peak frequency of approximately 6 Hz due of activity driven by backbone neurons (**Fig. 1B, top, Fig. 1C, left**). Meanwhile, the decrease in spiking interactions between inhibitory and backbone neurons simultaneously attenuated the faster PING-like theta oscillation associated with REM.

We further used a previously developed functional connectivity algorithm (72) (see **Methods**) to measure how functional connectivity shifts from NREM to REM for the modeled populations, based on spike timing relationships. We first quantified changes to mean pairwise functional connectivity (reported as a mean z-score; see **Methods**), either within the population of engram backbone neurons or within the population of LE neurons, as a function of outgoing backbone synapses’ strength (representing initial encoded memory strength). We systematically varied the strength of excitatory synapses from engram backbone (EB) to other excitatory neurons in the network, and separately, EB excitatory synapses targeting inhibitory interneurons (**Fig. 1E**). Strengthening synaptic connections from backbone neurons to other excitatory neurons (i.e., moving down the y-axis; **Fig. 1E**) increases mean functional connectivity in the network, while strengthening backbone neurons’ synapses on inhibitory interneurons (i.e., moving along the x-axis from left to right; **Fig. 1E**) decreases mean functional connectivity. However, these patterns differ between NREM and REM states. We observed that for backbone neurons’ connections with other backbone neurons, functional connectivity is generally weaker during NREM than REM, and does not vary as dramatically with incremental changes to synaptic strength. Conversely, mean functional connectivity between pairs of LE neurons is significantly higher during NREM than during REM. That naturally leads to a wider range of functional connectivity in REM than in NREM, consistent with recent data showing that firing rates are more divergent among hippocampal neurons in REM vs. NREM (73).

### Differential population-wide changes in pairwise functional connectivity between adjacent NREM and REM sleep bauds resemble experimental ones

We measured the population-level distribution of pairwise functional connectivity obtained from recordings from mouse hippocampal region CA1 undergoing single-trial contextual fear conditioning (CFC) (74, 75). Changes in NREM and REM sleep bouts’ functional connectivity were compared across 6h baseline and the first 6 h post-CFC (a time window critical for sleep-dependent memory consolidation (17, 74)), using methods identical for those used on model data (**Fig. 1E**). Namely, we measured changes of pairwise synaptic connectivity for the same pairs of recorded neurons in adjacent bouts of NREM and REM sleep (i.e., consecutive bouts of NREM and REM sleep that were closer that 10s apart). An example of such comparison for one of the animals is depicted in the left panel of **Fig. 2A**. We also analyzed these pairwise changes in functional connectivity for baseline and post-sham recordings in animals that were put to novel cage but did not receive an electric shock (74). We compared the baseline and post-experimental manipulation distributions for both cfc and sham animals, respectively. We observed that while in all the cases overall functional connectivity is shifted towards lower functional connectivity (i.e., smaller significance Z score values) during REM, the post-cfc animals show significant increase in number of neuronal pairs having higher functional connectivity from NREM to REM, as compared to their baseline (**Fig. 2A**, solid and dashed black lines; KS test comparison of baseline and post-cfc ***p=6*.*0102e-126***). Such a shift is not observed in sham animals (**Fig. 2A**, solid and dashed blue lines), with the two distributions showing minimally reverse trend (KS test comparison of baseline and post-sham ***p=0*.*0014***).

**Figure 2.**
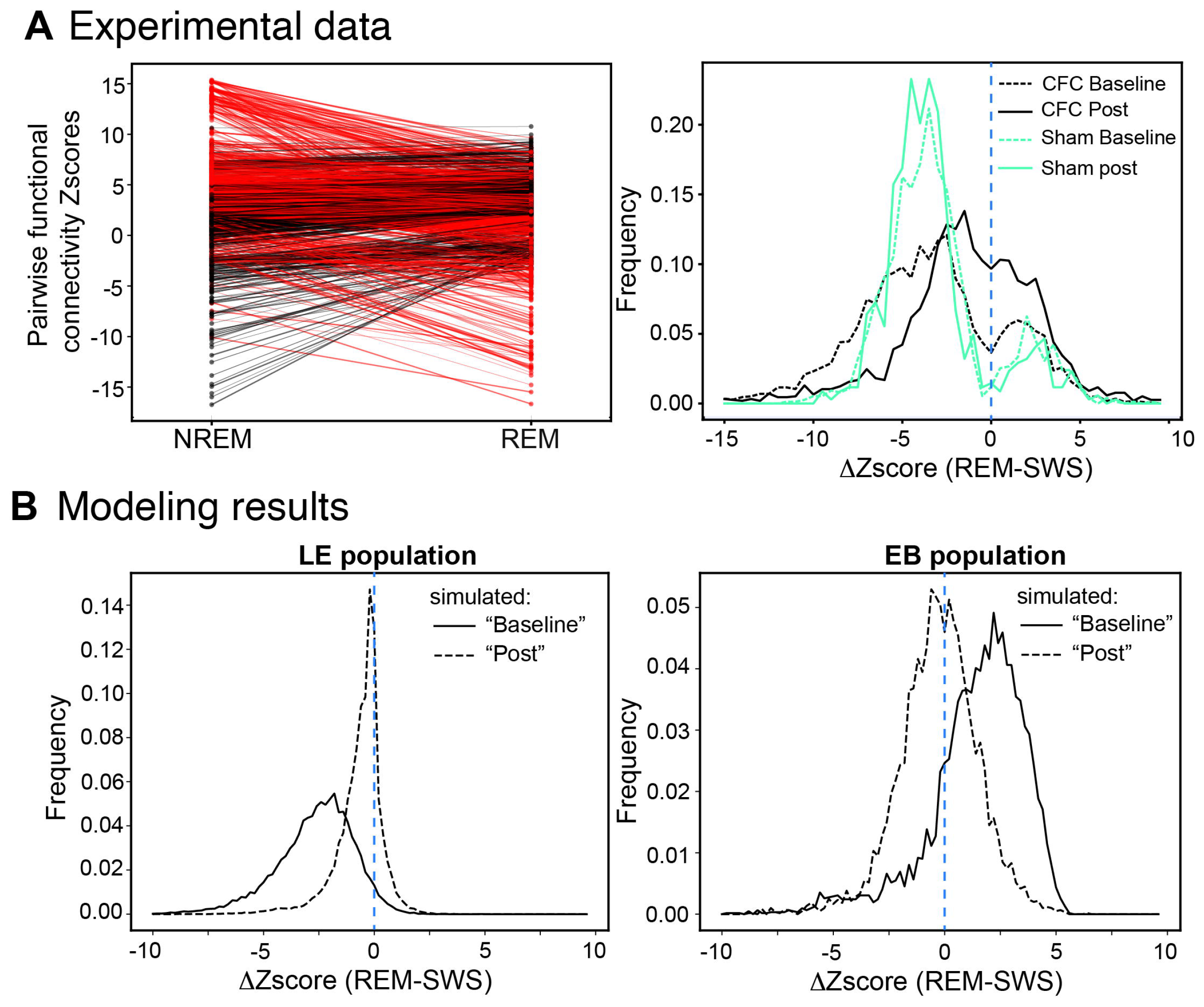
Changes in functional connectivity between NREM and REM – simulation and experiment. A) (Experiment) Change in functional connectivity (reported as change in z-scores) between adjacent NREM and REM sleep bouts (representative example of one animal – left panel; red – pairs with decreasing functional connectivity from NREM to REM, black - pairs with increasing functional connectivity from NREM to REM; pairs with |z|<2 in both REM and REM not shown). Right panel - baseline (dashed black line) and post-cfc (solid black line) changes for animals undergoing contextual fear conditioning, and similarly baseline (dashed blue line) and post-sham (solid blue line) changes for animals undergoing only sham training (i.e., no electric shock during novel cage presentation). REM generally weakens functional connectivity between following adjacent NREM. However for animals that underwent CFC exhibit significant increase in population of cell pairs that strengthen functional connectivity, as compared to baseline (K-S test comparing baseline and CFC p= 6.0102e-126). Such change is absent for sham animals, as they exhibit modest reverse trend (K-S test comparing baseline and sham p=0.0014). **B) (Modeling Results)** Change in pairwise functional connectivity (reported as change in z-scores) between simulated NREM and REM, for reduced (marked”B” on **Fig. 1E** and referred to as baseline) and full strength of excitatory coupling (marked”P” on **Fig. 1E** and referred to as post). Right panel - change in pairwise functional connectivity for baseline (dashed black line) and post (solid black line) for LE population. Left panel – similar change in pairwise functional connectivity for backbone population. REM prunes weaker connections associated here with LE population, shifting the distribution towards lower functional connectivity values (lower z-scores) as compared to NREM (center). Strong connections associated with backbone population can be however further strengthened.

We compared experimentally observed changes of pairwise functional connectivity with those observed in our model (**Fig. 2B**), and reported it, as above, as the difference in the Z-scores between the NREM and REM states. We compared these changes between simulated sleep states for two excitatory connectivity strength values for connections originating from EB population (as marked on **Fig. 1E**). We referred to reduced excitatory connectivity strength as simulated”baseline” (weak EB synaptic outputs - no new memory trace present; dashed lines on **Fig. 2B**) and to its standard strengthened value (i.e., new memory trace present) as simulated”post” (solid lines on **Fig. 2B)**. We measured changes in functional connectivity for those two cases, separately for less excitable (LE) population and engram backbone population. Both (LE and backbone) populations, when weakly coupled (i.e., at”baseline”) exhibit overall modest drop in pairwise functional connectivity from NREM to REM state (negative shift of baseline distributions, in both panels of **Fig. 2B**). In contrast, in strongly coupled case, LE and backbone populations exhibit dramatically different behavior. While pairwise functional connectivity of LE population is significantly reduced for LE population (**Fig. 2B, left panel**), the backbone population experiences significant shift towards higher significance values (**Fig. 2B, right panel**).

This result indicates that relatively weak connections, as those in LE population, during REM sleep get further weakened as compared to NREM. However, pairs exhibiting strongest pairwise functional connectivity, as those in EB population, get further strengthened.

### Differential roles NREM and REM states in recruiting heterogenous populations of LE neurons

We next investigated how NREM (low ACh, high *g*_*Ks*_) and REM (high ACh, low *g*_*Ks*_) differentially recruited LE neuron populations that receive different degrees of excitatory input from engram backbone neurons (**Fig. 3**). We divided the LE population into two groups - one receiving only weak coupling from backbone neurons, and a second where backbone neurons’ input is strengthened by a positive multiplier *M*_*ij*_ ∈ [1.0, 6.0] (**Fig. 3A**). We monitored population responses of LE neurons as a function of this backbone neuron input multiplier (*M* _*ij*_) during NREM- and REM-like states. During NREM, the LE population is disinhibited due to decreased firing rates among inhibitory interneurons (**Fig. 3B, top, red line**). This allowed both groups (i.e., those with and without the input multiplier) to fire in synchronous bursts with backbone neurons (**Fig. 3C, top**). This scenario provides optimal conditions for LE neuron populations to be recruited into the engram via STDP-mediated strengthening of inputs from engram backbone neurons (76) (**Fig. 3D, left**). **Figure 3D** depicts changes in mean synaptic strengths between three populations of excitatory cells (i.e., backbone neurons, LE neurons with *M*_*ij*_ =1, and LE neurons with *M*_*ij*_ =3), which are driven by STDP during post-learning sleep. During NREM, connections between backbone neurons and both LE groups are strengthened; strengthening also occurs to a lesser extent between neurons in the LE groups. In contrast, during REM, strong inhibitory input (**Fig. 3B, bottom, red line**) suppresses the activity of LE neurons with less excitatory input from backbone neurons (i.e., only LE neurons having high *M*_*ij*_ are active; **Fig. 3B, bottom, teal line**). The LE neurons that remain active receive a strong burst of excitation from backbone neurons and synchronize with them to form population bursts (**Fig. 3C, bottom**). These dynamics drive STDP-mediated strengthening of connections from backbone neurons to the few active LE neurons, while weakening their remaining excitatory connections to the rest of the LE neuron population (**Fig. 3D, right**).

**Figure 3.**
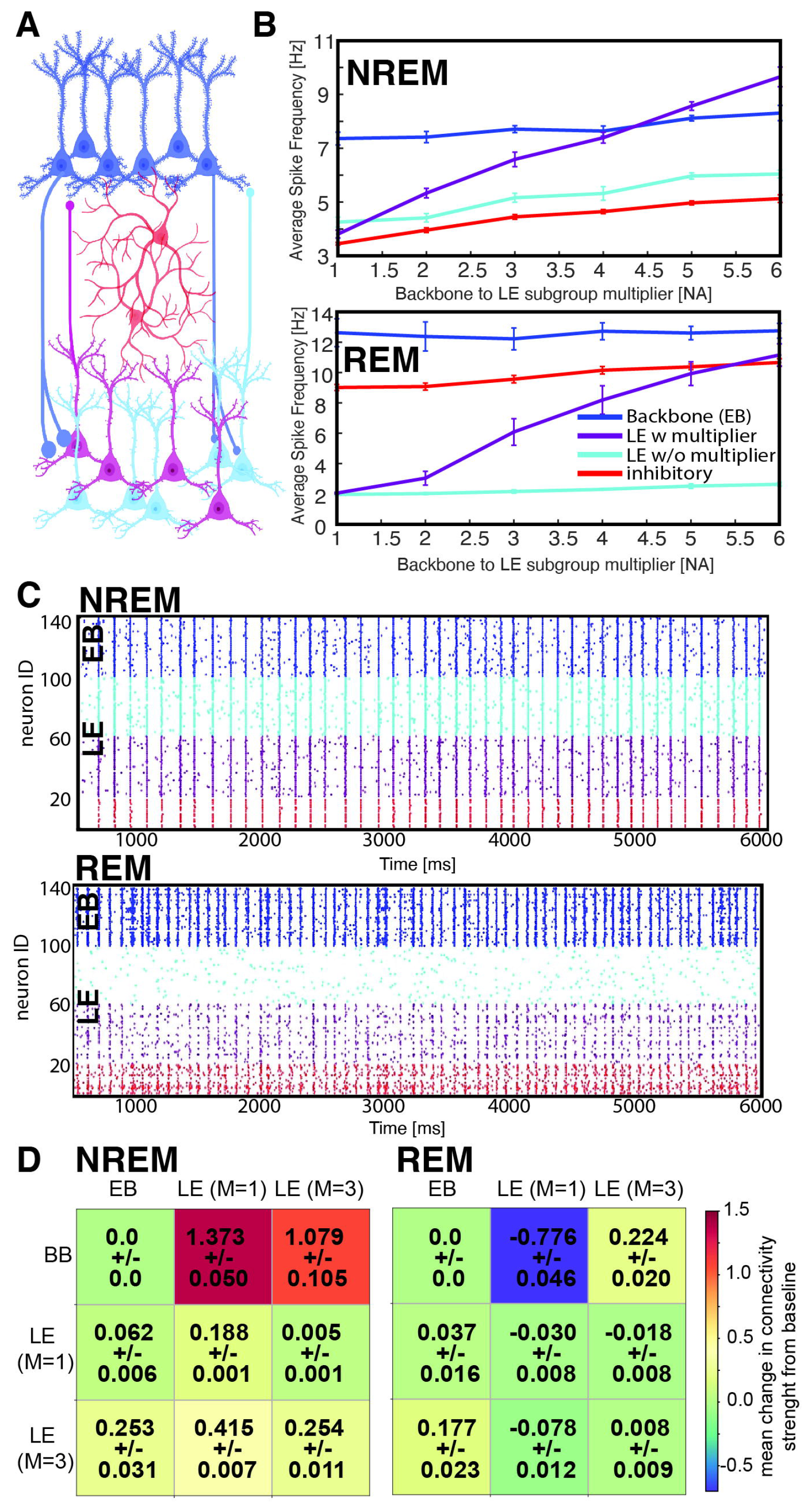
NREM and REM states differentially affect STDP-mediated incorporation of LE neurons into an engram. ***A)*** Schematic of the model network, consisting of memory-encoding backbone neurons (**EB, *dark blue***), inhibitory interneurons (***red***), and an LE group divided into two subpopulations - one receiving standard input from backbone neurons as shown in **Figure 1** (w/o multiplier; ***teal***) and one receiving variably enhanced input from the backbone population (w multiplier; ***violet***). ***B)*** Average firing frequency during NREM (**top**) and REM (**bottom**) for the four neuron populations, as a function of the LE subpopulation’s multiplier (*M*_*ij*_) value. ***C)*** Representative raster plot of firing during NREM (**top**) and REM (**bottom**) for the four neuron populations, using multiplier value of *M*_*ij*_ = 3. ***D)*** STDP-driven connection strength changes between and within the excitatory populations (i.e., backbone neurons and the two LE populations), for NREM (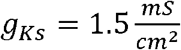, **left**) and REM (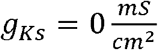, **right**). Plasticity between neurons within the backbone population is set to zero. Values indicate mean ± SEM across 4 simulation runs.

### Following encoding of multiple memories, NREM and REM differentially contribute to recruitment of network neurons into segregated engrams

Having characterized network activation and recruitment patterns as a function of ACh neuromodulation in NREM or REM, we next tested how sequential NREM and REM-like states affect memory consolidation (i.e., STDP-mediated recruitment of network LE neurons by engram backbone neurons) of two memories (**Fig. 4A**). The two memories were initially”encoded” in the network via activation of two independent backbone neuron populations, which had reciprocal cross-connections (shown in blue and green in **Fig. 4B**; neuron #s 100 and above). Within the network, initial connections from each engram backbone were assigned randomly to LE neurons; thus the number of connections each LE neuron received from either backbone varied. For visualization purposes, we divided the LE population into four quartiles (**Fig. 4B** and **BB**): 1) those receiving the strongest input at baseline (i.e., most connections) from backbone neurons encoding memory 1 and also the weakest input (i.e., least connections at baseline) from backbone neurons encoding memory 2 (blue with gray background), 2) those receiving the strongest input from memory 2 and the weakest from memory 1 (green with gray background), 3) those receiving relatively weak connections from both memory 1 and memory 2 backbone neurons (pink), 4) those receiving strong connections from both memories (violet). To model reduced overlap of the synaptic projections from memory engrams activated by backbone neurons we reduced the depolarizing input 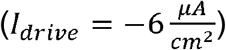 to both, pink and violet groups in LE population, so that they fire only randomly and sparsely, and are not recruited into the engram. Here, in addition, the green and blue LE groups receive initially 20% stronger input from memory 1 and memory 2 backbones, respectively (ω_*ij*_ (*t*=0) =1.2; **see Methods**). We, however, tested how the degree of this biasing of the connections (**Supplementary Fig. S1**) affects both, the recruitment (referred to as”activation”) and segregation (see **Methods**) of the LE populations into the engrams of memory 1 and memory 2 during NREM and REM, respectively. We observed that both processes are robust and occur even without modification of connection strengths (**Supplementary Fig. S1**).

**Figure 4.**
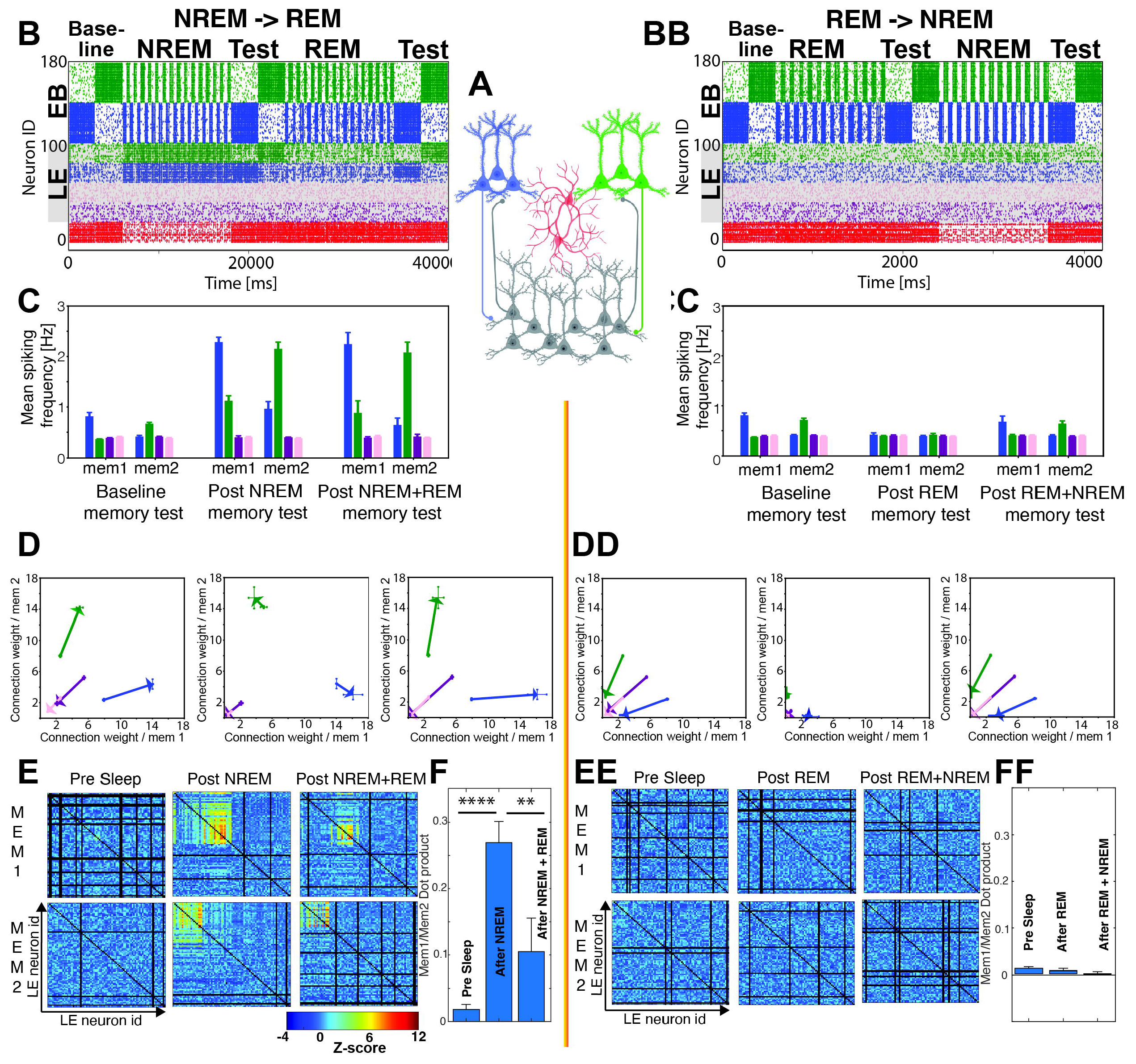
Differential roles of NREM and REM, and benefits of wake→NREM→REM sequences, during consolidation of multiple memories. Network simulations are run with stereotypic wake→NREM→REM sleep architecture (**left column; *B-F***) or with reversed wake→REM→NREM architecture (**right column; *BB-FF***). ***A)*** Schematic of the model network, where the backbone neuron population (EB) consists of two separate groups encoding memory (***blue***) and memory 2 (***green***). ***B)*** Representative raster plot of network activity, where retrieval/activation patterns of LE neurons are tested in response to memory 1 and memory 2 backbone population activation before sleep (pre-sleep), after NREM, and after NREM+REM. Tests occur in high Ach 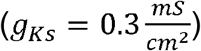, with continuous input from one of the two backbone populations, and no synaptic plasticity. Backbone neurons’ firing is not shaded (neuron IDs 100-180); firing of all LE neurons is shaded in gray (neuron IDs 20-80). LE neurons receiving stronger input (i.e., more connections at baseline) from the memory 1 backbone population are shown in blue; those receiving stronger input (i.e., more connections at baseline) from the memory 2 backbone population only are shown in green. Pink and violet LE neurons indicate populations receiving weak (i.e., least connections from both backbones at baseline) and strong input from both backbone populations (i.e., most connections from both backbones at baseline), respectively; these groups receive lower constant current (*I*_*drive*_) than green and blue LE groups, which leads to reduced, more random, firing patterns. ***D)*** Changes of synaptic connectivity between the two memory backbone populations and the four LE groups, between following timepoint tests: baseline to post NREM (left), post NREM to post REM (center), baseline to post NREM+REM (right). X-axis denotes mean connection weight from memory 1 backbone to LE layer cells; the Y-axis denotes mean connection weight (bottom) from memory 2 backbone to LE layer cells. The vectors demark the change of the mean connections strengths between the afore mentioned timepoints (see **Fig. S1**) ***E)*** Pairwise functional connectivity among LE neurons during memory tests. Colors indicate the statistical significance (bootstrapped Z-score) of connections during pre-sleep, post-NREM, and post-NREM+REM tests for a representative simulation. ***F)*** Overlap of functional connectivity patterns among LE neurons during reactivation of memory 1 vs. memory 2 backbone populations, during pre-sleep, post-NREM, and post-NREM+REM tests. Values indicate mean ± SEM for 4 simulation runs; ^****^ and ^**^ indicate *p* = 0.004 for pre-sleep vs. post-NREM, and *p* = 0.033 for post-NREM vs. post-NREM+REM, respectively. ***BB - FF)*** Panels showing functional connectivity changes (similar to ***B – G***) for a reversed-architecture sleep cycle (i.e., REM followed by NREM). REM— NREM state ordering leads to weakened connectivity between memory-encoding backbone neuron populations and LE neurons, leading to failed recruitment of LE neurons into memory traces during sleep.

We monitored the effects of the two sleep states on memory consolidation during”test” epochs, which loosely represent wake-state memory retrieval. These tests - where m-current is low (i.e., ACh release is high) and backbone neuron populations representing one of the two memories are reactivated - occurred at baseline, before sleep consolidation (baseline), and then after each sleep of the two sleep states (**Fig. 4B**). Within NREM and REM states, STDP was allowed to take place at excitatory synapses - between backbone and LE neurons, and between LE neurons. During this time, the engram backbone populations’ reactivation was driven via periodic depolarizing current injection at theta frequency. We observed that during pre-sleep testing, blue and green LE neuron populations fire only sparsely in response to activation of either the memory 1 or memory 2 backbone population. During subsequent NREM sleep (**Fig. 4B**), both the blue and green LE populations become more highly active via disinhibition, and their firing is largely independent of the backbone populations’ activation pattern. That leads, via STDP, to indiscriminate potentiation of connections between the two backbone populations and LE neurons. This in turn leads to increased and overlapping activation of the LE populations during post-NREM testing, where LE neuron activation can be driven by either of the backbone populations. When REM follows NREM, inhibition to LE neurons becomes significantly stronger, causing them to fire more sparsely, with firing locked to the backbone population they receive the strongest input from (i.e., green LE neurons become phase locked to memory 2 backbone neurons, and blue LE neurons become phase-locked with memory 1 backbone neurons). These dynamics cause STDP-mediated weakening of connections from the backbone population that were weaker prior to initial testing. Thus, during subsequent post-REM testing, there is a larger segregation between the activated LE populations - where LE neurons activated downstream of the original memory 1 and memory 2 backbone populations become largely non-overlapping.

To quantify recruitment of LE neurons into memory representations we characterized changes to four observables during the three memory test epochs (i.e., pre-sleep baseline, post-NREM, and post-NREM→REM): 1) changes in firing frequency of the respective LE neuron groups (**Fig. 4C**); 2) changes in synaptic connectivity between memory 1 and memory 2 backbone populations and these LE neuron groups (**Fig. 4D and** Fig. S2), and 3) changes in patterns of functional connectivity between the LE neurons (**Fig. 4E, F**). The firing frequency of LE neurons preferentially coupled to memory backbones dramatically increases (relative to baseline) during the post-NREM testing phase. Firing in the two strongly-coupled LE groups tends to be higher during reactivation of the memory backbone that they receive the strongest connections from, but they have significant activation in response to reactivation of either memory. After subsequent REM, the difference in firing rates between the two strongly-backbone-coupled LE neuron groups increases, leading to greater discrimination based on their respective firing during activation of the two memory traces (**Fig. 4C**). These state-driven changes in firing rates during testing were related to STDP-driven changes in excitatory synapse connection strength between memory 1 and memory 2 backbone neuron populations and the LE neuronal subgroups. **Supplementary Fig. S2 (diagonal)** depicts sample distributions of all connection strengths from memory1 (x-axis) and memory 2 (y axis) backbone populations to all LE neuron subgroups during pre-sleep, post-NREM and post REM testing epochs. The pink and violet groups lie on the diagonal, indicating similar excitatory input from the two memory backbone populations. After NREM, both green and blue LE neuron groups show similarly strengthened connections along both axes. Due to only sparse random activation during NREM (caused by reduced input as compared to other two groups), the pink and violet LE neuron groups are synaptically decoupled from the two backbones. The”x” marks mean values for each group, respectively. The mean change in connection strength between pre-sleep and NREM test epochs (averaged over four simulation runs) is depicted on **Fig. 4D** and off-diagonal terms of **Supplementary Fig. S2**, with colored vectors corresponding to changes in mean excitatory connection weight across NREM (**Fig. 4D; left panel; Supplementary Fig. S2**). After subsequent REM, the blue LE neuron group now receives excitatory input predominantly from the memory 1 backbone, whereas the green group receives input predominantly from memory 2 backbone. The mean change in connection strength between post NREM and post NREM→REM tests is shown in **Fig. 4D, center plot**. Ultimately, the NREM→REM sequence results in separated populations of LE neurons with strongly potentiated synapses from only one of the two memory backbone populations (**Fig. 4D, right plot**).

We also analyzed patterns of functional connectivity within the LE neuron population, using the functional connectivity algorithm described for **Fig. 1E**, above (72). We used this metric to characterize pairwise coactivation patterns between LE neurons when memory 1 and memory 2 backbone populations were activated separately during testing pre-sleep, post-NREM and post-NREM→REM (**Fig. 4E**). While no significant functional connectivity between LE neuron pairs was observed during the pre-sleep test period, strong connectivity was observed both between, and within, green and blue LE neuron groups after NREM. After the NREM→REM sequence, functional connectivity between the two groups during memory testing was reduced significantly, but intra-group connectivity was preserved. This suggests that NREM followed by REM drives segregation of recruited LE neurons into separate populations corresponding to different memory traces. To further address this, we quantified the degree of overlap between the LE neuron populations with significant functional connectivity to memory 1 or memory 2 backbone populations (**Fig. 4F**). This overlap is zero during the pre-sleep test, increases significantly after NREM alone, and decreases after the NREM→REM sequence.

Finally, we measured the population-level distribution of pairwise functional connectivity among all LE neurons (**Supplementary Fig. S3**) for both, memory one (**Supplementary Fig. S3A, top**) and memory 2 (**Supplementary Fig. S3A, bottom**). After NREM sleep alone, most of the connections have undergone strengthening, which causes the distribution to be strongly skewed towards positive values **(Supplementary Fig. S3A, left panels**). In comparison, after the following REM, there is a strong tendency for connections to be weaker, although a small fraction of connections have strengthened significantly even compared to NREM alone (**Supplementary Fig. S3A, center panels**). Thus, while synaptic weakening and loss of functional connectivity is the predominant effect of REM, some functional connections get stronger during this time. Finally, we also compared changes in the functional connectivity between the baseline and the post-NREM+REM test **Supplementary Fig. S3A, right panels**). While most functional connections among the LE population were not significantly changed, a fraction of connections were strengthened, corresponding to LE neurons that were selectively recruited into either of the two memories’ engrams (**Fig. 4E**).

To test the robustness of these findings, the duration of the depolarizing current pulses used for reactivation during NREM and REM, and the temporal separation of reactivation for the two backbone populations, were systematically varied, as shown in **Supplementary Figs. S4** and **S5**. The duration of reactivation had little impact on the either recruitment of the LE neurons into engrams (referred to as”activation”, see **Methods**), or the segregation of the memories within the LE population (referred to as”segregation”, see **Methods**; **Supplementary Fig. S4**). However, the temporal separation of reactivation between the two backbone populations has an impact on both properties, with longer delays between reactivation of memory 1 and memory 2 backbones leading to decreases in both properties. However, lengthening the total duration over which reactivation occurred (and thus the total number of reactivations) could compensate for both disrupted strengthening of connections to LE neurons, and segregations of the two memories’ connections to the LE population, when delays in reactivation between memory 1 and 2 were longer (**Supplementary Fig. S5**). Together, these data suggest that after NREM→REM sequences, engrams corresponding to memory 1 and memory 2 become both stronger (activating larger populations of neurons) and more distinct from one another (with less population overlap).

We have also tested the effects of activation of the subpopulation of LE neurons that receives highest number of connections from both memory backbones (i.e., population marked in violet). Namely we varied the difference in the *I*_*Drive*_ between this population and the other two LE populations (demarked as blue and green on **Fig. 4B**). Not surprisingly we observe that with strong activation of the pink population the segregation capability of the network is reduced significantly, whereas the activation of the LE network remains high (**Supplementary Fig. S6)**. This suggests that if the two memories have a large initial overlap in the network (i.e., they both activate a large common subpopulation of activated LE cells) than they cannot be segregated and instead form a common engram.

### Sequential NREM→REM state ordering is essential for memory consolidation

Because wake→NREM→REM sleep state sequences are ubiquitous (and wake→REM→NREM sequences are not typically observed), we were curious how reversing the sequence would impact the outcome of consolidation. We tested the effects of the REM-like sleep state preceding the NREM-like state (**Fig. 4BB-FF; right column**). In this scenario, we observe diametrically different memory consolidation dynamics (**Fig. 4BB**). Increased synaptic inhibition during REM suppresses activity among LE neurons and causes depression of memory 1 and memory 2 backbone neuron connections to LE neurons. Synaptic inputs from the backbone populations, once weakened, can no longer drive burst firing among LE neurons during subsequent NREM engram reactivation. This eliminates recruitment of the LE neuron population into engrams during NREM. This is evident at the level of LE population firing rates during memory testing; firing rate responses evoked by memory 1 or memory 2 backbone populations decrease after initial REM, and minimally recover after subsequent NREM (**Fig. 4CC**). Similarly, the strength of backbone populations’ input connections to LE neurons decreases after initial REM (**Fig. 4DD**), effectively disconnecting LE neurons from memory backbone populations. Functional connectivity clusters do not form in this scenario (**Fig. 4EE**) and the population overlap between memory 1 and memory 2 activation remains zero (**Fig. 4FF**), indicating a complete lack of recruitment of LE neurons into the memory engrams. There are also no observable changes in functional connectivity at population level (**Fig. 4EE, Supplementary Fig. S3B**) due to the complete lack of LE neuron recruitment. Thus, if the typical NREM→REM sequence is reversed (to REM→NREM), the end result is a lack of engram enlargement – i.e., a failure of memory consolidation.

### Multiple sleep sequences drive progressive improvement in memory segregation

Sleep, across species, occurs in repeating cycles of NREM and subsequent REM. These cycles occur either intermittently throughout the day, or in the case of humans and some other species, in rapid succession throughout a longer, grouped sleep period. An unanswered question is whether, and why, it might be beneficial to have multiple NREM→REM cycles, rather than a single, massed period of NREM and REM. To address this, we ran two additional sets of simulations. In the first, we divided the sleep period into 4 repeating NREM→REM cycles, with each cycle followed by a memory test in which memory 1 and memory 2 backbone populations were separately activated as described above (**Fig. 5A-B**). In the second, memory was tested after a single NREM→REM sequence, with the total duration of both NREM and REM matching the combined length of the sleep states (i.e., from all 4 cycles) from the first simulation (**Fig. 5C-D**). For the test periods interspersed between the multiple, shorter cycles (**Fig. 5B**), the green and blue LE neuron groups activate preferentially with their respective memory backbone neurons, maintaining minimal cross-activation. After the longer-duration, single NREM→REM cycle, there is significant co-activation of these LE neuron populations during testing (**Fig. 5C-D**). We quantified this effect by defining parameters that we refer to as”activation” and”segregation” (**Fig. 5C**; see **Methods**). Activation (black line) indicates the mean firing frequency of green and blue LE neuron groups when the corresponding backbone neuron population is activated during each test phase. Segregation (red line) is the normalized difference in mean firing frequency between the green and blue LE neuron groups during their respective testing phases. We observe that both activation and separation increase steadily as a function of the number NREM→REM sleep cycles. The activation for non-participating LE neuron groups (pink and violet) does not change throughout the sleep cycles (green line). However, the outcome of a longer-duration, single NREM→REM cycle was qualitatively and quantitatively different. Activation of the two LE neuron groups was significantly greater than after cycling sleep, but their segregation was decreased. This means that while the two neuron groups are more strongly recruited into the memory traces than after multiple, shorter NREM→REM cycles, the crosstalk between the two recruited populations is much larger. This is because during extended NREM, connections from the memory 1 and 2 backbone populations to green and blue LE neuron groups become highly potentiated, as do the connections between these two groups. Once green-blue”crosstalk” connections become sufficiently strong (i.e., during extended NREM), subsequent REM dynamics can cause simultaneous, consistent co-activation of the two LE populations, preventing typical REM-mediated depression and pruning of these connections (and thus memory segregation). Taken together, these data suggest that in certain circumstances (e.g., when consolidating more than one memory in a network simultaneously), multiple NREM→REM cycles can be more effective than a single, longer-duration cycle for keeping engram populations segregated.

**Figure 5.**
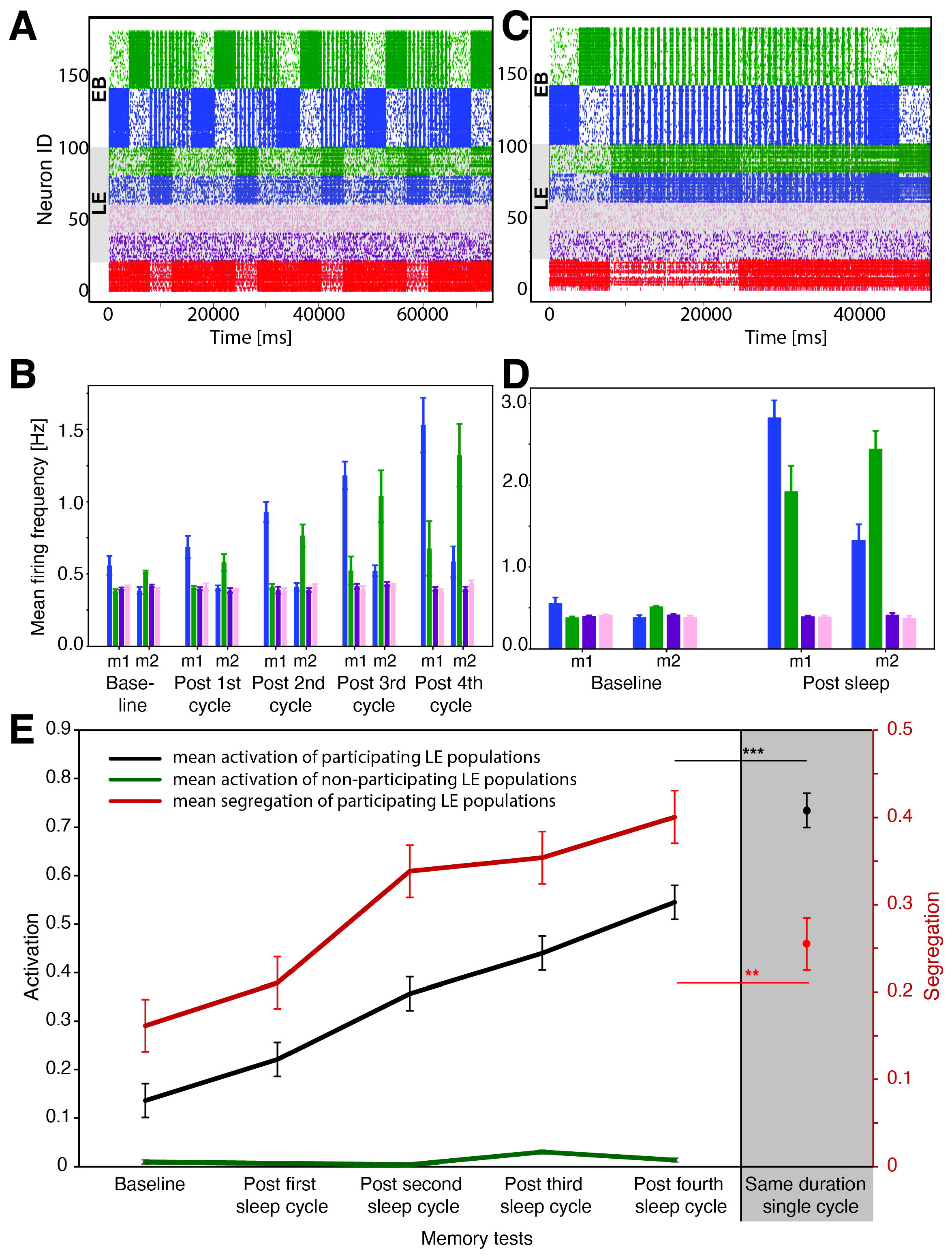
Benefits for consolidation of multiple NREM→REM cycles. ***A)*** Representative raster plot over four NREM→REM cycles. Memory retrieval tests were performed as described for **Figure 3**, after every NREM+REM sequence. ***B)*** Mean firing responses within LE groups during activation of memory 1 or memory 2 backbone neuron (EB) populations. ***C)*** Representative raster plot depicting a single, extended NREM→REM cycle (i.e., of the same total duration as the four cycles in ***A***). Retrieval tests were performed before and after the NREM+REM sequence. ***D)*** firing responses within LE groups during retrieval-associated activation of memory 1 or memory 2 backbone neuron populations. ***E)*** Changes in activity profiles (activation and segregation; see **Methods**) for LE groups after multiple cycles vs. a single sleep cycle of the same total duration (**gray area**). Multiple sleep cycles result in greater segregation of engram representations corresponding to different memories. Values indicate mean ± SEM for 4 simulation runs; ^***^ and ^**^ indicate *p* = 0.004 and *p* = 0.038, respectively, for activation and segregation.

## Discussion

Vertebrate sleep cycles, while variable in length, consist of repeating wake→NREM→REM transitions; this ordering of sleep states is nearly universal (77). However, the specific roles of NREM and REM in sleep functions such as memory consolidation (3, 78), and the functional significance of this ubiquitous sleep state ordering, are completely unknown. Sleep (taken as a whole) has been proposed to play various roles in regulating synaptic changes in the context of memory consolidation (4), from reactivation of neurons and networks engaged by prior learning (21, 79, 80), to regulation of excitatory-inhibitory balance (9, 14, 81), to homeostatic downscaling of synapses (82, 83). Here we have used a simplified biophysical model - in which state-associated network dynamics are regulated primarily through changes in a muscarinic ACh receptor-modulated current – to show that NREM and REM sleep may play distinct, complementary, and necessary roles in sleep-dependent memory consolidation.

We show that changing ACh level during NREM and REM directly modulates neuronal excitability via this pathway, and that this in turn affects the excitatory/inhibitory balance of the network. During NREM, low ACh permits m-current suppression of inhibitory neurons’ firing, leading to network disinhibition. This scenario allows for STDP-driven recruitment of initially less-active excitatory neurons into memory engrams, driven by learning-activated backbone neuron populations (**Figs. 1** and **3**). In contrast, during REM, m-current is suppressed by high ACh levels, and the resulting elevated network inhibition (**Fig. 1**), suppression of activity in most of the less-active excitatory neurons (sparing those most strongly driven by backbone populations), and competitive pruning of newly formed memory representations (**Fig. 3**). This mechanism captures changes in firing statistics observed experimentally in CA1 in the hours following learning (as opposed to sham situation, **Fig. 2**), where REM generally exhibits lower functional connectivity as compared to NREM, except for limited populations of cells that have their functional interactions strengthened.

NREM→REM sequential cycling plays especially critical role when consolidating multiple memories (**Fig. 4**), as it allows for engram expansion while keeping the representations of consolidated memories separated from one another. Within the model, reversing the sequence (i.e., REM→NREM) leads to a failure of memory trace expansion (**Fig. 4**) and replacing multiple NREM→REM cycles with one prolonged sleep cycle leads to a failure of memory segregation when two memories are being consolidated simultaneously (**Fig. 5**). Together, these findings provide a theoretical basis for why seemingly-ubiquitous sleep architecture and transitions may be critical for the memory-processing aspects of sleep function.

Critically, several of our present modeling results are supported by recent *in vivo* data. For example, preventing the normal decrease in ACh signaling during NREM sleep disrupts memory consolidation, in both human subjects (57) and mice (14). Numerous recent findings support the idea that memory traces may expand, though disinhibition and subsequent synaptic potentiation, during NREM. In regions such as the hippocampal dentate gyrus, suppression of inhibitory transmission during sleep seems to be an essential component of memory processing (14). Available data suggests that reactivation (or sequential replay) of memory traces occurs preferentially (and more broadly throughout the network) during NREM as a correlate of active memory consolidation (16, 17). Finally, there is accumulating evidence of synaptic strengthening occurring preferentially during post-learning NREM sleep. The analyses shown in **Fig. 2** illustrate that, as is true for our model network data, functional connectivity between CA1 neurons is stronger during post-learning NREM vs. post-learning REM, except for smaller number of pairwise interactions that show competitive strengthening. *In vivo* electrophysiological recordings have demonstrated baseline (19, 84) or learning-driven (18, 20) firing rate increases that occur selectively across NREM bouts. Finally, *in vivo* recordings have shown that potentiation of thalamocortical synapses occurs selectively during NREM bouts (85), and *in vivo* imaging data have illustrated dendritic spine formation during post-learning NREM (11). There are also strong data to support the notion that REM facilitates inhibition-driven pruning of memory traces. Recent *in vivo* imaging data have shown that in neocortex, interneuron-mediated suppression of pyramidal cell activity is higher during REM sleep compared with NREM sleep (9). The idea that excitatory synapses are selectively pruned during REM (but not NREM) is supported by *in vivo* electrophysiological recordings from neocortex and hippocampus (where principal cell firing rates decline selectively across REM bouts) (84, 86) and *in vivo* imaging studies of neocortex (where dendritic spines are lost preferentially during REM) (10).

While the model network used here was intentionally designed with a simple and generic structure, its features are applicable to neocortical and hippocampal networks that are studied in the context of memory consolidation *in vivo*. For example, the mechanism underlying the differential functions we observe in NREM vs. REM hinges on the role inhibitory interneurons play in sculpturing memory representations (**Fig. 4**), and the state-specific regulation of interneuron activity by ACh. Recurrent feedback inhibition is thought to promote competition amongst principal neurons throughout the brain. Reliable spiking of interneurons is observed in the hippocampus and other structures after spontaneous pyramidal cell firing (87), or after experimental (i.e., optogenetic) activation of one to several pyramidal cells (88-90). On the other hand, optogenetic silencing of principal neurons can lead to strong disinhibition of neighboring principal cells (91, 92). This mechanism is known to be important in regions like the dentate gyrus, where interneuron populations are known to gate the population of principal neurons activated during memory consolidation (14) and recall (93). ACh preferentially activates interneuron populations in the hippocampus (53, 54) and neocortex (94-96); this selective sensitivity is mediated by both nicotinic and muscarinic receptors (55, 97). Because ACh release from medial septal inputs to hippocampus, and by basal forebrain inputs to neocortex, is higher during wake and REM vs. NREM sleep (98-101), this indicates that as is true in our model, these networks are gated by interneurons during REM and disinhibited during NREM. Thus, while the model does not incorporate more circuit-specific sleep-state features (i.e. localized modulation by other neuromodulator systems, interneuron subpopulation-specific effects, or specific network oscillations)(3, 4, 102-104), it recapitulates this more fundamental aspect of brain state regulation.

Behavioral studies in both human subjects and animal models have aimed to identify precise roles of NREM vs. REM sleep in specific aspects of memory consolidation, to whether the two states have differential roles. The importance of NREM sleep for consolidating many forms of memory (including declarative and procedural memory) has been well established, based on studies either disrupting NREM behaviorally, targeting state-specific features of NREM, or enhancing NREM-specific network activity patterns (14, 16, 18, 23, 105). Our modeling results - showing that engram populations generally expand through a combination of reactivation and synaptic potentiation during NREM sleep - are consistent with these data. On the other hand, only some types of memories rely on post-learning REM sleep (i.e., they are interrupted by REM-targeted sleep disruption), while many others appear to require NREM but not REM for long-term storage. In rodent behavioral studies, tasks disrupted by post-learning REM sleep deprivation tend to be more complex in nature – i.e., those in which multiple associations (e.g. place-reward contingencies) must be made to ensure proper task performance (5). Effects of REM sleep-targeted deprivation studies in human subjects have been less clear, with many forms of memory consolidation (particularly declarative memory) proving robust to this manipulation. However, some sleep-dependent forms of consolidation are disrupted by prevention of REM, including studies where multiple associations (e.g., of different visual cues with an aversive shock vs. safety) must be made in more complex scenarios (106). A converse approach, which has been used successfully to address REM’s cognitive functions in human subjects, is identification features of behavioral tasks where benefits are correlated with post-learning REM sleep. In one such study, participants were trained on two competing sensory discrimination tasks in rapid succession, followed by a nap opportunity. Only those who engaged in a post-learning nap containing REM showed perceptual improvement for both tasks, without interference between them (62). Finally, a recent study used administration of ACh receptor antagonists to block ACh transmission during post-learning REM sleep in human subjects, and found subtle deficits in consolidation. Intriguingly, however, subjects in the study were trained on two separate memory tasks prior to the intervention (107). Together, these data support the idea that REM sleep with high ACh signaling could provide an opportunity to segregate competing memories from one another, to improve performance on complex tasks and/or prevent interference between memories.

REM seems to facilitate other cognitive functions, which may not immediately appear related to a role in memory storage. Intriguingly, data from human subjects indicates that following a period of REM (but not NREM), participants show increased creative thinking, have greater ability to form new associations, and are more likely to have restructured learned information (108, 109). These features are not incompatible with the”pruning” aspects of REM described here, insofar as elimination of weak or irrelevant connections throughout a network may increase its capacity for forming new connections upon receiving new inputs. Experimental results from both humans and animals show that the elevated neural inhibition observed during REM protects memories from interference to permit continual learning (110).

Our present data provide a set of testable hypotheses about the differential roles that NREM and REM sleep play in sleep-dependent memory consolidation. They suggest specific neural network-level mechanisms associated with particular behavioral effects of NREM and REM in the consolidation process, wherein NREM drives expansion of memory traces and subsequent REM maintains optimal segregation of traces when multiple memories are being encoded. The predictions indicate a critical role of REM based pattern separation during storage of multiple correlated (in terms of time and content) memories. Finally, our data suggest an essential biological function for the seemingly ubiquitous wake→NREM→REM cycling phenomenon, which is present in nearly all vertebrate species.

## Methods

### 1. Neuron model

Excitatory and inhibitory neurons were modeled using a conductance-based Hodgkin-Huxley formalism (111, 112). The time-dependent voltage *V*_i_ of an individual neuron is given by the master equation:

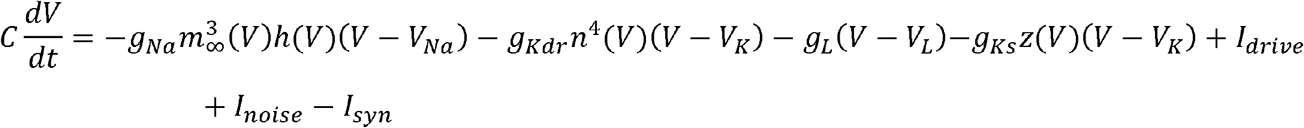

with the first four currents on the right-hand side representing a fast inward Na^+^ current, a delayed rectifier K^+^ current, a membrane leak current, and a slow outward (hyperpolarizing) K^+^ current representative of muscarinic M_1_ receptor activation (m-current). The m-current conductance, *g*_*Ks*_, serves as a proxy for the state-dependent ACh level, with ACh reducing activation of this current. During NREM sleep, when ACh levels are low, m-current is high 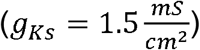; conversely, during REM sleep, when ACh concentration is high, m-current is blocked (*g*_*Ks*_ → 0). On a cellular level, *g*_*Ks*_ conductance modulates membrane excitability, with low *g*_*Ks*_ (high ACh) yielding Type 1 excitability, whereas high *g*_*Ks*_ (low ACh) yielding Type 2 excitability. Type 1 excitability is characterized by low firing frequencies in the absence of excitatory input, but a high frequency gain in response to input (i.e., a steep current-frequency [i-f] curve), and advance-only firing phase responses to brief stimulation pulse (i.e., phase response curve [PRC]). Type 2 excitability has a threshold in firing frequency onset, a shallow frequency gain function (i-f curve), and a biphasic PRC *(111)*. The parameters in the above equation (except for driving current [*I*_*drive*_], described below) are the same for both excitatory and inhibitory interneurons, and are provided in **Table 1**.

**Table 1.** Cellular parameters of modeled excitatory and inhibitory neurons.

| Parameter | Value |
| --- | --- |
| $C$ | $1 \frac{\mu F}{cm^2}$ |
| $g_{Na}$ | $24 \frac{mS}{cm^2}$ |
| $g_{Kdr}$ | $3 \frac{mS}{cm^2}$ |
| $g_{KS}$ | $0 - 1.5 \frac{mS}{cm^2}$ |
| $g_L$ | $0.02 \frac{mS}{cm^2}$ |
| $V_{Na}$ | 55 mV |
| $V_K$ | -90 mV |
| $V_L$ | -60 mV |

The gating variables, i.e., sodium inactivation, *h*(*v*), potassium activation, *n*(*v*), and m-current activation, *z*(*v*), are given by:

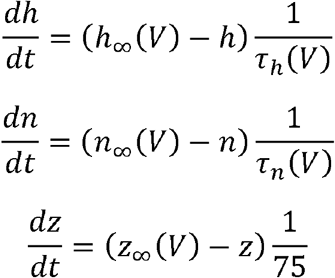

Where *h* _∞_(*V*), *n* _∞_(*V*), *z* _∞_ (*V*), *m* _∞_(*V*), τ _*h*_ (*V*), τ_*n*_ (*V*) and are defined as:

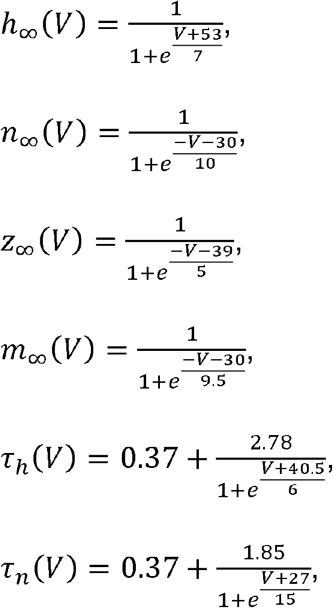

where the time constants are expressed in [ms] and voltages in [mV]. External driving current,*I* _*drive*_ is a parameter used to set the excitability of neurons. For excitatory neurons, 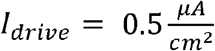 unless otherwise specified, while for inhibitory neurons 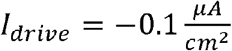. The difference in excitability between the two populations ensures that inhibitory neurons fire only in response to excitatory input, providing natural competition between excitatory populations. Random input, *I*_*noise*_, representing input from other external modalities, is modelled by a low-probability excitatory current influx to each neuron. Every millisecond there is a 0.002% chance that a given neuron will receive an 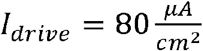 input with duration of 1 ms. This results in spontaneous firing with mean frequency of 2 Hz for all neurons in the network. Finally, *I*_*syn*_ represents synaptic input from other network neurons, and is described in detail below.

### 2. Network architecture

#### Single-memory network

The network encoding a single memory trace (**Figs. 1** - **3**) is composed of 120 excitatory neurons and 20 inhibitory neurons. Excitatory neurons provide sparse and random input to the network, with 10% connectivity to both excitatory and inhibitory neurons. Inhibitory neurons provide input to 50% of neighboring excitatory and inhibitory neurons within a specified connectivity radius. Excitatory neurons are divided into two groups, the encoded memory’s initial backbone (engram backbone EB) population (40 neurons) and less activated (LE) neurons (80 neurons). Backbone neurons represent the initial memory trace encoded during learning, receiving less inhibition than LE neurons, and having higher-efficacy, fixed synaptic weights between one another. LE neurons fire less due to decreased interconnectivity and increased inhibition (see section below). Excitatory synaptic inputs targeting LE neurons (i.e., from both engram backbone and LE neurons) undergo spike timing-dependent plasticity as described below.

#### Two-memory network

The two-memory network (**Figs. 4** and **5**; **Supplementary Figs. S1**-**S6**) is composed of 160 excitatory and 20 inhibitory neurons. The difference in number of excitatory neurons stems from creating a second engram backbone, as these neurons are divided evenly into three groups – 80 LE neurons, 40 backbone neurons corresponding to memory 1, and 40 neurons corresponding to memory 2. The two memory-encoding backbone populations have no synaptic connections with one another, but provide random, sparse input to both inhibitory and LE neuron populations. Note that the two engram backbones are activated sequentially, thus the maximal current generated by EB population remains largely the same. Other two-memory network parameters are identical to the single-memory network.

Since input connections from each backbone population to LE neurons are random, the number of connections each LE neuron receives from either backbone population varies. We divided the LE neuron population into four quartiles (see supplementary Fig. S2): (1) **stronger input from memory 1 (i.e**., **neurons receiving most connections at baseline from mem 1 backbone; denoted as blue in the LE population – gray background on Figs. 4, 5)** – this quartile receives relatively strong input from the memory 1 backbone population and relatively weak input from memory 2, (2) **stronger input from memory 2 (i.e**., **neurons receiving most connections at baseline from mem 2 backbone; denoted as green in the LE population – gray background)** - this quartile receives relatively strong input from the memory 2 backbone population and relatively weak input from memory 1, (3) **weak input from both of the memories (i.e**., **neurons receiving least connections at baseline from both backbones; denoted as pink in the LE population)**, and (4) **strong input from both of the memories (i.e**., **neurons receiving most connections at baseline from both backbones; denoted as violet in the LE population)**. In most of the simulations, the last two populations (i.e., pink and violet) have sparse and random firing (driven by noise), as they receive lower *I*_*drive*_ as compared to other LE groups. We have tested the effects of activating the population of LE cells receiving the largest number of synapses from both memory backbones (violet group), and as predicted we observe that strong activation of these neurons hampers segregation of the two memories (**Supplementary Fig. S6**)

### 3. Synaptic communication and plasticity

When each modeled neuron’s membrane voltage exceeds preset threshold (5 mV), an action potential is recorded, and the presynaptic neuron activates inhibitory or excitatory postsynaptic current to all its postsynaptic targets. The postsynaptic currents,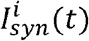, received by an (i-th) cell are modeled as a linear combination of fast and slow excitatory and inhibitory currents.

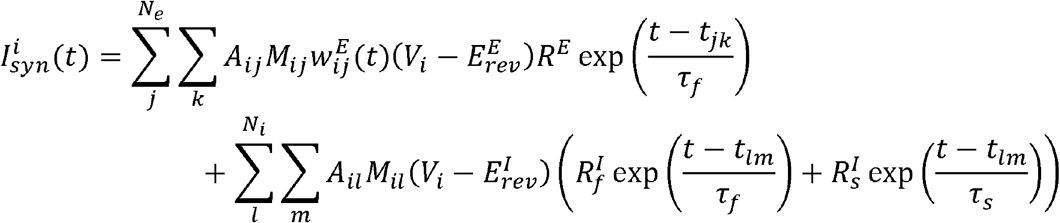

Where *t*_*j*k_ is the timing of *k*-th spike on *j*-th neuron; 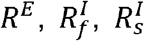 denote relative amplitudes of excitatory, fast inhibitory and slow inhibitory currents, respectively (see **Table 2**). Time constants τ_f_ =0.5*ms* and τ_f_ =50 *ms* define the time course of fast and slow post-synaptic currents, respectively. 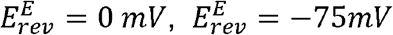 are reversal potentials for excitatory and inhibitory currents, respectively. Synaptic multiplier *M*_*ij*_ differentiates synaptic efficacy of inputs from specific neuron populations to others (see **Table 3**). The term 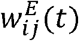 indicates time-dependent synaptic weight changes driven by STDP between excitatory *i*-th and *j*-th neurons.

**Table 2.** Relative synaptic efficacies between the neuron types.

| Synapse Type | Postsynaptic target neuron type |  |
| --- | --- | --- |
|  | Excitatory | Inhibitory |
| $R^e$ (excitatory) | 0.15 | 0.08 |
| $R_f^i$ (fast inhibitory) | 0 | 0.15 |
| $R_s^i$ (slow inhibitory) | 0.05 | 0 |

**Table 3.** Synaptic group multipliers. Intra-backbone connections are stronger than those between the LE neurons; the LE population receives stronger inhibitory input than backbone neurons.

| $M_{ij}$ | | Postsynaptic | | |
| --- | --- | --- | --- | --- |
|  |  | Backbone neurons | LE neurons | Inhibitory interneurons |
| **Presynaptic** | Backbone neurons | 2.5 | 1 | 1 |
|  | LE neurons | 1 | 1 | 1 |
|  | Inhibitory neurons | 0.5 | 3.5 | 1 |

**Table 4.** Population-specific magnitude of synaptic plasticity. Plasticity of the synapses emanating from LE population is significantly slower than that of synapses originating from backbone population. Only excitatory-to-excitatory synapses undergo plastic changes.

| $P_x$ | | Postsynaptic | | |
| --- | --- | --- | --- | --- |
|  |  | Backbone neurons | LE neurons | Inhibitory interneurons |
| **Presynaptic** | Backbone neurons | 0 | 1 | 0 |
|  | LE neurons | 0.3 | 0.3 | 0 |
|  | Inhibitory interneurons | 0 | 0 | 0 |

*A*_*ij*_ denotes terms in the adjacency matrix. 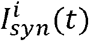 is represented in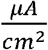.

For simulations that included STDP (**Figs. 3-5**), the change in synaptic strength between postsynaptic neuron *i* and presynaptic neuron *j* was given by:

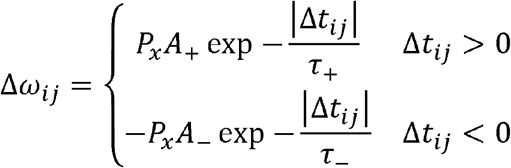

where Δ*t*_*ij*_ is the difference between the most recent spike time of postsynaptic neuron *i* and presynaptic neuron j; A_+_ and A_-_and *A*_*-*_ are the maximum rates of change (excluding *P*_x_) with values A_+_ =0.07 and A_-_ =0.025. τ_+_ and τ_-_ are the time constants, τ_+_ =14 ms and τ_-_ =34 ms. The overall effect is plasticity that favors potentiation between neurons with more synchronous firing (Δ*t < ∼25ms*) and favors depression for longer inter-spike intervals. This leads to overall synaptic depression in a randomly and sparsely firing network, and to synaptic potentiation in a highly active network. *P*_x_ is a network-wide controlled parameter for changing the rate of plasticity between specific neuronal groups (see **Table S4**). These parameters are set in a way that only excitatory-to-excitatory synapses undergo plastic changes, and the speed at which these changes occur is slower at inputs from the LE population than from backbone populations. Across intervals of STDP-associated plasticity, synaptic weight *W*_*ij*_ for each connection is constrained to the interval [0, *W*_max_] where *W*_max_ =5. Initially synaptic weights were set to ω_*ij*_ (*t*=0) =1 for most excitatory-to-excitatory connections, except for those of the memory 1 backbone targeting the LE population that receives strong connections from memory 1 only (i.e., blue group), where ω_*ij*_ (*t* =0) =1.2. Similarly, ω_*ij*_ (*t*= 0) =1.2 for the memory 2 backbone synapses targeting the LE population that receives strong connections from memory 2 only (i.e., green group). We have tested the effects of this connection bias (**Fig. S1**) and observe that it has only minimal effect on both activation and segregation of the LE populations.

### 4. Memory testing: activity, synaptic strength, and functional connectivity analyses

Average mean firing rates (**Figs. 1, 4 - 5**) were calculated for the LE population overall, or separately for each subgroup of the LE population described above. These measures were used to calculate mean strength and segregation of LE engrams’ representation of memory 1 and memory 2. Activation of memory 1 vs. memory 2 (*A*_*½*_) is defined as 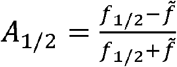, where *f*1/2 is the mean firing frequency during test activation of memory1 or memory 2 backbone neuron populations (i.e., memory testing), and 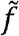 is the mean frequency of the non-activating LE neuron populations (groups denoted by pink and purple on the raster plot on **Figs. 4** and **5**). It measures the relative activation of the blue and green groups of LE neurons above background. Segregation of memory 1 from memory 2 (*S*_*½*_) is defined as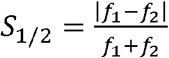, where *f*_1_ (*f*_2_) is the mean firing frequency of memory1 (memory 2). It measures differential activation of the two LE representations during each respective memory test.

Mean synaptic strengths of inputs from memory 1 and memory 2 backbone populations (x-axis and y-axis, respectively, **demarked as ‘x’ on Supplementary Fig. S2 diagonal**) were calculated for each LE subgroup defined during every test phase. Changes in mean synaptic strength between tests was calculated as a vector (**Fig 4D, DD and Fig S2** (off-diagonal)), by positioning the tail at the onset coordinates and head at the final coordinates. Synaptic strength change vectors were calculated between pre-sleep and post-NREM sleep tests (or post-REM sleep tests for reverse-order simulations), between post-NREM and post-REM tests, and between pre-sleep and post-NREM+REM tests (or post-REM+NREM tests for reverse-order simulations).

To quantify the pairwise functional connectivity between LE neurons (**Fig. 1E, Fig. 2B, Fig. 4E** and **EE**) and from experimental recordings (**Fig. 2A**), a metric based on the average minimal distance (AMD) was used (113). The pairwise AMD between two neurons (*i, j*) was given by the mean difference of the time of each spike, *k* in spike train of the *i*-th neuron, *S*_*i*_, to the most recent preceding spike in the spike train of *j*-th neuron, *S*_*j*_. This was calculated as:

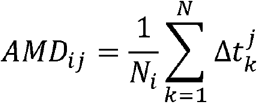

where *N*_*i*_ was the number of spikes in spike train *S*_*i*_ and 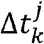 was the temporal distance between *k*-th spike in *S*_*i*_ to the nearest spike in *S*_*j*_ (113). To quantify the magnitude of temporal locking between the pair of neurons (i.e., the magnitude of their functional connectivity), a z-score was calculated, using *L* as the length of the inter-spike interval of spike train *S*_*i*_. The first and second moments (*μ*_1_ and *μ*_2_, respectively) for the spike train *S*_*i*_ are given by 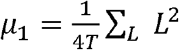 and 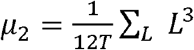, where *T* is the total time of the *S*_*i*_ -th spike train (in ms). These were used to derive mean and standard deviation of the minimal distance with respect to *S*_*i*_, where the mean μ = μ_1_, the first moment, and the standard deviation 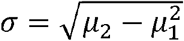 (113). The Z-score is given by 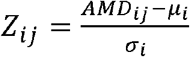 (see (17, 72)). Negative z-scores imply that the neurons fire in less coincident fasion than random firing would instigate, indicating inhibitory interactions. Large z-scores indicate tight temporal coincidence.

To establish separation of two memories (**Figs. 3F** and **FF**), the dot product of functional connectivity matrixes was calculated between testing periods when memory 1 vs. memory 2 backbone populations were reactivated. Data were averaged over 4 randomly-initialized simulations. Histograms of functional connectivity changes (**Fig. 4A** and **Supplemental Figure S6**) were calculated using the pairwise z-score distributions of every connection pair across compared test phases.

### 5. *In vivo* hippocampal recordings, and contextual fear conditioning (CFC)

All animal husbandry and experimental procedures were approved by the University of Michigan Institutional Animal Care and Use Committee. Male C57BL/6J mice between 2 and 6 months (*n* = 4) were implanted with custom built driveable headstages with two bundles of stereotrodes implanted in bilateral CA1, and EMG electrodes to monitor nuchal muscle activity *(74, 114)*. The signals from each of the stereotrode recording sites were split into local field potential (0.5-200 Hz) data, which were used to evaluate behavioral states, and spike data (200 Hz-8 kHz), which were used for functional connectivity analysis. Following post-operative recovery, mice underwent habituation to daily handling and tethered recording as stereotrodes were lowered into CA1 to obtain stable recordings. After establishing stable single-neuron recordings, each mouse underwent single-trial CFC (placement into a novel environmental context, followed 2.5 min later by a 2-s, 0.75 mA foot shock) starting at lights on (i.e., ZT 0), after which they were returned to their home cage for *ad lib* sleep with continued recording. Spike data from individual neurons was discriminated offline using standard methods (consistent waveform shape and amplitude on the two stereotrode wires, relative cluster position of spike waveforms in principle component space, ISI ≥ 1 ms) *(74, 75, 115-117)*.

Functional connectivity (**Fig. 2A**) of neurons recorded from individual animals was calculated separately for each adjacent bout pair of NREM and REM sleep (REM onset needed to be <10s following the NREM offset), using the AMD method described above. To offset length discrepancies between NREM and REM bouts affecting the statistical comparison, we analyzed only the final segment of NREM bout matching time duration of the paired (following) REM. The analysis was performed on both CFC and sham animals (i.e. no electrical shock during presentation of novel cage); 6h baseline was compared to 6h post experimental manipulation (i.e., CFC or sham).

## Acknowledgements

This work was supported by NIH MH135565 (SJA, MZ), NIH NS118440 (SJA), NSF-1757574 (AS), and University of Michigan Department of Physics (MS).

