## Supplemental material for "Neuromodulation via muscarinic acetylcholine pathway can facilitate distinct, complementary, and sequential roles for NREM and REM states during sleep-dependent memory consolidation"

**Supplemental Information**

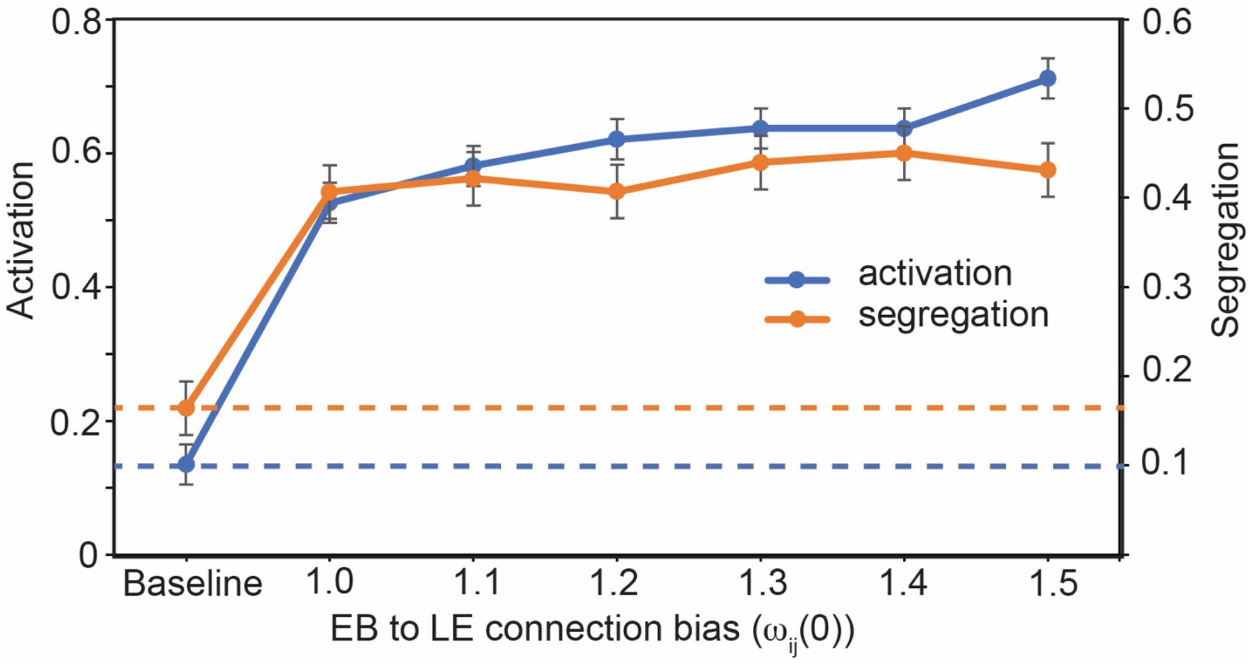


**Figure S1. Activation and segregation of memory 1 and memory 2 in LE layer as a function of connection multiplier, $\boldsymbol{\omega_{ij}(t=0)}$ (see methods).** When $\omega_{ij}(t=0) = 1.0$ all LE groups have the same weight initially and the only difference between assingned LE subpopulations is a random fluctuation in the number of connections at baseline, from EB layer (as decribed in methods). Activity at baseline is measured before the sleep epochs occur. The multiplier $\omega_{ij}(t=0)$ does not affect significantly neighter activation or segregation of recruited LE neurons into memory 1 and memory 2.

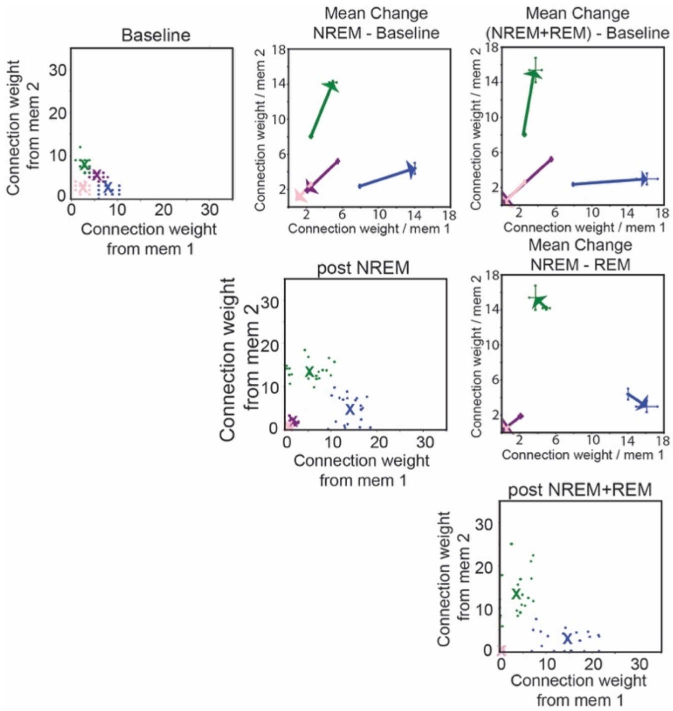


**Figure S2. Connection strength changes during simulated sleep memory consolidation.** **DIAGONAL:** representative connection maps from backbones of both memories to individual neurons in LE layer. Each dot represents a total connection strength (i.e., sum of synaptic efficacies of neurons belonging to one of the backbones and targeting given LE neuron) from backbone of mem 1 (X-axis) and mem 2 (Y-axis) to individual LE neurons. The connectivity is set at random. The whole population of LE layer at baseline is divided into 4 quartiles (and remains the same for the rest of the simulation): LE neurons receiving stronger input (i.e., more connections) from the memory 1 backbone population are shown in blue; those receiving stronger input (i.e., more connections) from the memory 2 backbone population only are shown in green. ***Pink*** and ***violet*** LE neurons indicate populations receiving weak (i.e., least connections from both backbones) and strong input from both backbone populations (i.e., most connections from both backbones), respectively. During most of the simulations (except **Fig. S5**) these groups receive lower constant current ($I_{drive}$) than ***green*** and ***blue*** LE groups, which leads to reduced, more random, firing patterns. **Top left**: representative map obtained at baseline. **Center**: representative map obtained post NREM. **Bottom right:** representative connection map obtained post NREM+REM. X - denotes mean connection strength for the given population.
**OFF-DIGONAL:** Change of mean connection strength between following timepoint tests: baseline to post NREM, post NREM to post REM, baseline to post NREM+REM. Values indicate mean values of 4 simulation runs.

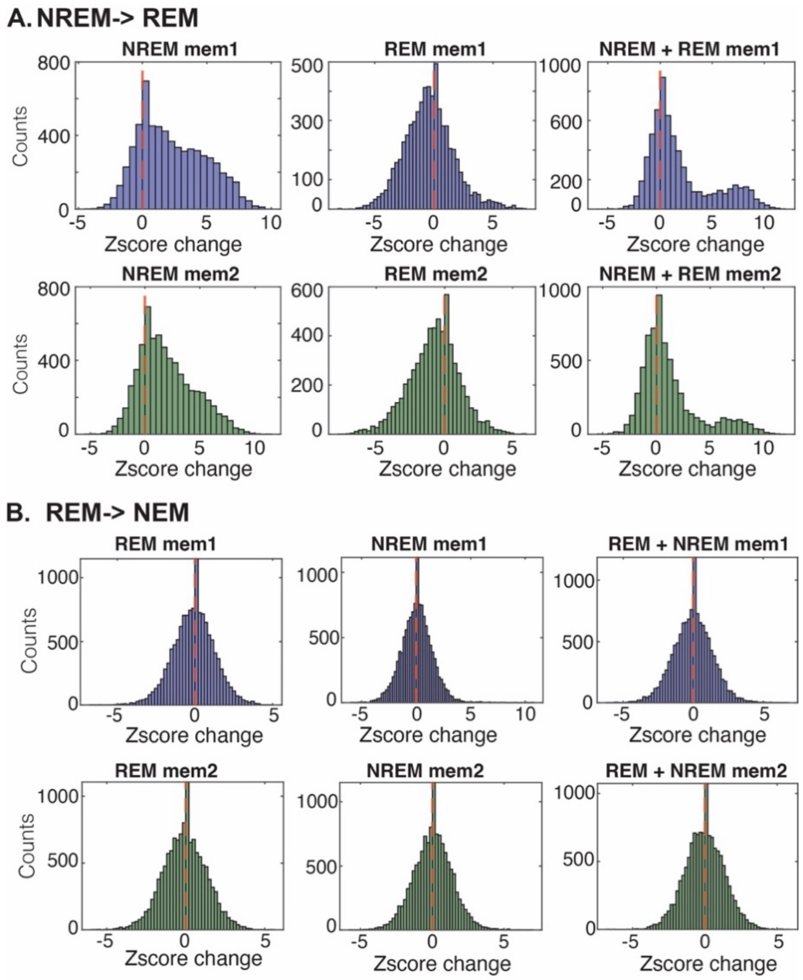


**Figure S3.** ***A)*** **Histograms of pairwise functional connectivity changes between excitatory neurons after NREM, after REM, and after a NREM+REM sleep cycle.** NREM sleep indiscriminately recruits LE neurons into the engram by strengthening connections from backbone neurons to LE neuron populations. This results in a shift of the functional connectivity distributions towards stronger connections (higher z-scores) within the LE neuron population (left). REM prunes these connections, leaving/strengthening only the strongest ones, shifting the distribution towards lower functional connectivity values (lower z-scores) and compared to NREM (center). When full sleep cycle (REM+REM) is compared with baseline (right) overall most of the connections don't change significantly except for relatively small group of connections that underlie recruitment of the LE cells into the engram.

***B)*** **Histograms of pairwise functional connectivity changes between excitatory neurons after REM, after NREM, and after a REM+NREM sleep cycle, when the sleep cycle is reversed (i.e., REM precedes NREM).** Because REM prunes backbone of LE connections before LE cells could be recruited into the engram, there are no significant shifts in the distributions of the connections.

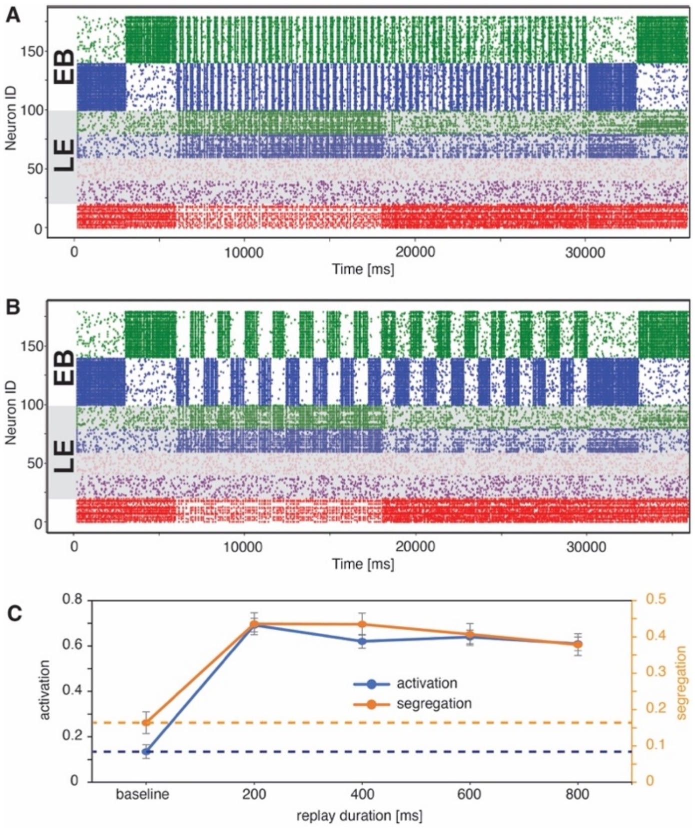


**Figure S4. Activation and segregation of the recruited LE representations as a function of length of reactivation bauds of memory 1 and memory 2 backbones.** Reactivation bouts were varied between 200-900 ms. ***A)*** Sample rasterfor 200 ms reactivation bauds. ***B)*** Sample raster for 800 ms reactivation baud. ***C)*** Calculation of activation and segregation (see **Methods**) as a function of length of reactivation bauds. Both activation and segregation are largely independent of reactivation baud length. Results in ***C*** averaged over 5 simulation runs.

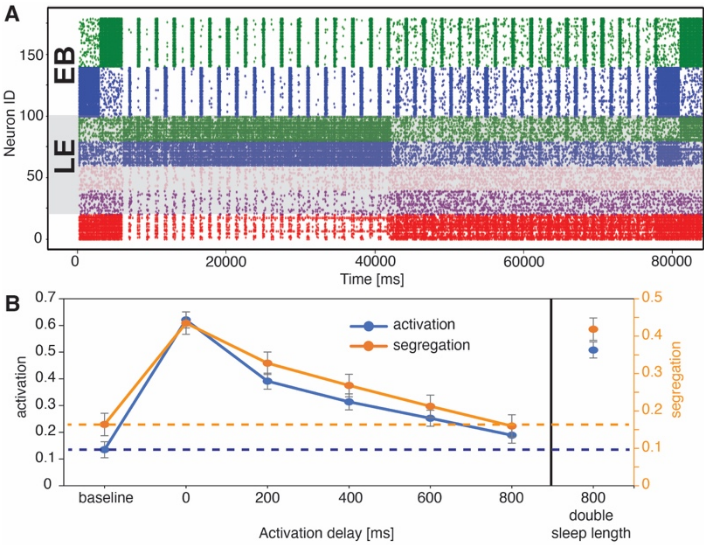


**Figure S5. Activation and separation of the recruited LE representations as a function of activation delay between reactivation bauds of memory 1 and memory 2 backbones.** The reactivation bout duration is set to 200 ms. The reactivation delay (i.e., dead space between two consecutive reactivations) is varied between 0-800 ms. ***A)*** Sample raster of simulation with an activation delay of 200 ms. ***B)*** Activation and segregation of LE representations (see **Methods**) as a function of length of reactivation delay. Both activation and segregation decrease as a function of activation delayed. However, when the simulation length is controlled for total reactivation time (which decreases as a function of reactivation delay), both functions recover. This indicates that total reactivation time controls activation and segregation magnitude rather than reactivation delay. Values in ***B*** indicate mean of 5 simulation runs.

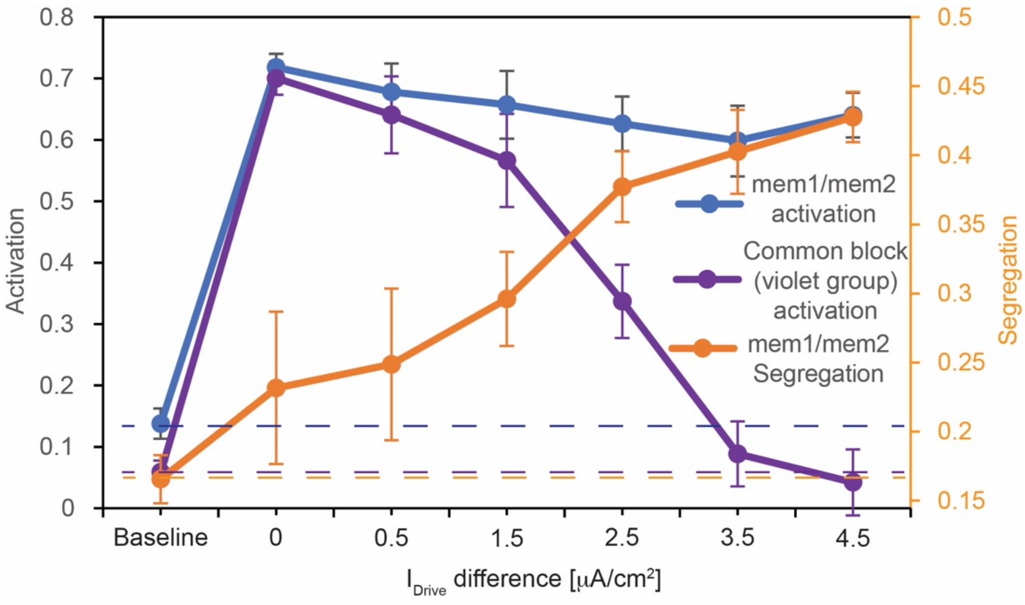


**Figure S6. Activation and segregation of LE neurons belonging to mem1/mem2 as a function of activation of neurons shared by both memories (i.e., LE sub-population that has largest number of connections with backbones of both memories; group denoted as violet on Figs 4, 5 and Figs. S2, S4, S5).** To activate this sub-population of LE cells, we changed its $I_{Drive}$value from that of the other two (i.e., ***green*** and ***blue***) LE populations; invariably, $I_{Drive}$=-6$\frac{\mu A}{cm^2}$ for the population having the least connections with the backbones of the two memories (i.e, group dentoted as pink **on Figs 4, 5 and Figs. S2, S4, S5**). The x-axis denotes difference in $I_{Drive}$ between the violet and green/blue LE population. When the $I_{Drive}$ is the same for all the groups the ***common subpopulation*** activates strongly (violet line), as do the both memories, mem1 and mem 2 (***blue*** line). The segregation (orange line) is however impeded. As the pink LE population is progressively inactivated (larger values of $I_{Drive}$difference) the segregation returns to normal levels. This intuitively indicates that if the memories share large common engram population the memories can not be segregated and a single consolidated engram forms.
